# Influence of sustained cognitive loading on finger circulatory and thermoperceptual responsiveness to localized cooling

**DOI:** 10.1101/2025.02.15.638420

**Authors:** Maaike I. Moes, Antonis Elia, Ola Eiken, Michail E. Keramidas

**Author notes:** **Corresponding Author:** Michail E. Keramidas, Division of Environmental Physiology, Department of Physiology and Pharmacology, Karolinska Institute, Berzelius väg 13, 17 65, Solna, Sweden.

## Abstract

Our aim was to examine whether finger vasomotor and thermoperceptual responses to local cooling would be modulated by sustained cognitive loading. To this end, finger temperature, circulatory (i.e., cutaneous vascular conductance, CVC) and perceptual responses were monitored, in twelve healthy men, during and after a 30-min hand immersion in 8°C water, performed either immediately after a 60-min continual execution of a cognitive task battery (cognitive→cold trial), or during the simultaneous performance of the cognitive task (cognitive+cold trial). Subjects’ responses were compared with those obtained in a control cold-provocation trial, wherein they watched an emotionally-neutral documentary. The cognitive task temporary enhanced the perceived levels of mental effort and fatigue in both intervention trials. In the cognitive→cold trial, the cold-induced reduction in finger CVC and increase in mean arterial pressure were blunted (*P* < 0.01), and the thermal discomfort was alleviated (*P* = 0.05). In the cognitive+cold trial, no intertrial differences were noted during the cold-water immersion phase (*P* ≥ 0.28), but the finger CVC was enhanced during the last part of the rewarming phase (*P* = 0.05). Present findings, therefore, demonstrate that *(i)* in moderately mentally-fatigued individuals, finger cold-induced vasoconstriction is transiently attenuated, and thermal discomfort is mitigated, and *(ii)* superimposition of cognitive loading on cold stress does not alter finger vasoreactivity or thermosensitivity during cooling, but facilitates finger reperfusion following cooling.

## INTRODUCTION

Local cold stress prompts an adrenergically-mediated increase in the tone of cutaneous vessels, limiting heat loss to the environment (for reviews see Refs. 1, 2). In acral skin regions (e.g., fingers, toes), however, the initial vasoconstriction is often interrupted by episodic elevations in blood flow and temperature (aka cold-induced vasodilation, CIVD; 3), which alleviate thermal discomfort and pain (4). The precise mechanism underlying the CIVD response is unclear (for reviews see Refs. 5, 6); yet the anatomical structure of the acral skin where CIVD predominantly occurs suggests that the arteriovenous anastomoses (AVAs), which are densely innervated by adrenergic axons (7), may play a major role (8).

Aside from thermal influences, cutaneous vasomotion is also subject to non-thermal factors, such as arousal stimuli (e.g., see Refs. 9-12). Specifically, in thermoneutral conditions, a brief period of mental stress, provoked by, for instance, a series of fast-paced arithmetic calculations, appears to promptly enhance the sympathetic outflow to the cutaneous vascular beds (13–15), increasing blood flow in the non-acral (e.g., forearm) skin arterioles (16–22), and, conversely, reducing flow in the AVA-dense acral-skin areas (e.g., finger; 9, 21, 23-28). The acral-skin vasoconstrictive response may also prevail when a cognitive stress task is superimposed on localized cooling: the cold-induced vasoconstriction is aggravated, and the incidence and magnitude of the CIVD response are suppressed (29–31). The acral-skin constrictor responsiveness to mental stress is, however, not a consistent finding, and its prevalence may to a large extent depend on the state of autonomic nervous activity preceding the mental-stress application. Previous works have thus yielded evidence that, in conditions wherein the basal sympathetic vasomotor tone is already high (e.g., during whole-body cooling, or in patients with Raynaud’s disease), the subsequent administration of an acute cognitive stressor may not augment the cold-evoked constriction, but, rather, induce a transient blood-flow elevation in the acral-skin arterioles (23, 25, 27, 32–34). It appears that information on such stressor interactions, is as yet limited to the impact of short-term (i.e., ≤10 min) mental stress on cutaneous vasomotor responsiveness to cold, whereas the influence of extended periods of cognitive loading, produced by engagement in a mental task requiring sustained levels of effort and attention, presumably leading to some degrees of fatigue (cf. Ref. 35), remains unclear.

The present study, therefore, aimed to examine whether finger vasoreactivity and thermosensitivity to direct, localized cooling would be modulated by cognitive loading associated with the prolonged performance of a mentally demanding task. For this purpose, we used a within-subject design, during which a cohort of healthy male individuals underwent a hand cold (8°C-water) provocation trial, either immediately after (*trial 1*) or during (*trial 2*) the repeated execution of a cognitive task battery requiring higher-order function resources. We hypothesized that, compared to in a control cold-provocation trial wherein the cognitive stimulation was maintained relatively minimal (*trial 3*), finger vasoconstriction would be attenuated when the cold stress was preceded by cognitive loading, whereas, by contrast, it would be augmented when the thermal and cognitive stressors co-existed.

## METHODS

### Ethics approval

The experimental protocol was approved by the Regional Human Ethics Committee of Stockholm (Ref. No.: 2022-06823-01, 2024-05578-02) and conformed to the standards set by the *Declaration of Helsinki*, except for registration in a database. Prior to participation, subjects were informed in detail about the experimental procedure and gave their written consent.

### Subjects

Thirteen healthy, right-handed men were recruited. One subject, however, exhibited, during his first experimental session, signs of cold-induced hypotension (36) upon immersing the hand into 8°C water; the trial was terminated immediately, and the subject withdrew from the study. Twelve subjects were thus included; their characteristics are summarized in **TABLE 1**. They were normotensive, non-smokers, and had no history of cold injury. All subjects were university students, physically active on a recreational basis (i.e., 1-3 exercise sessions·week^-1^), and did not participate habitually in cold-water activities (e.g., winter bathing or swimming). They were instructed to abstain from alcohol and heavy exercise for at least 24-h before each trial, to refrain from caffeine during the testing day, and to maintain their sleeping (≥7 hours·night^-1^), eating, and exercise routines throughout the study period; subjects’ adherence to these instructions was confirmed by individual logbooks.

**TABLE 1.**
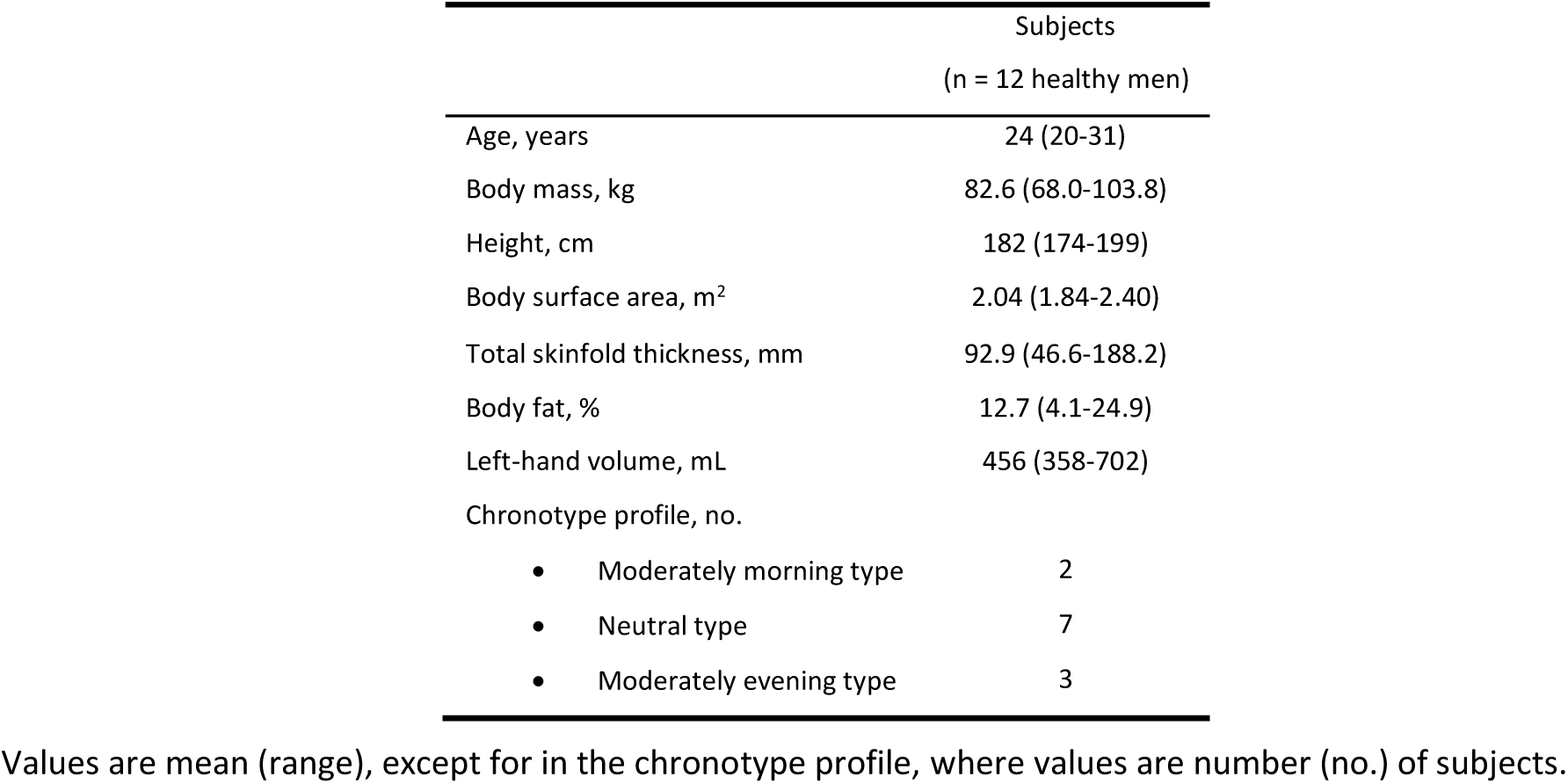
Subjects’ characteristics.

### Study design and testing protocol

Subjects initially attended a preliminary session, wherein they underwent a medical examination and were familiarized with the testing procedure. They were thus asked to immerse their left hand in 8°C water for <5 min, and to perform 5-6 times, interspersed by a >5-min interval, the cognitive task battery (*see below*). Anthropometric measures were also conducted: (i) body mass and height were measured, and the body surface area was calculated (37), (ii) the skinfold thickness was measured at seven right-sided locations, and the body fat percentage was calculated (38), (iii) the volume of the left (i.e., tested) hand was estimated using a water displacement method, and (iv) the individual chronotype profile was assessed with the morning-eveningness questionnaire (39).

Approximately a week after the preliminary session, each subject participated in three experimental sessions (days), being performed in a counterbalanced, Latin-square fashion, and separated by ≥48 h. During each session, subjects underwent a local cold provocation trial, consisting of immersing the left hand in 8°C water for 30 min, followed by a 15-min spontaneous rewarming. The three sessions protocols were:

1. The provocation trial was preceded by a 1-h cognitive task (i.e., *cognitive→cold* trial).
2. The provocation trial was performed concomitantly with the cognitive task (i.e., *cognitive+cold* trial).
3. The provocation trial was performed without any cognitive task (i.e., *control* trial).

The testing protocol is depicted in **FIGURE 1**. In all trials, subjects remained quietly in an upright sitting position, with both hands supported at the level of the heart. Throughout the trials, the right (i.e., non-immersed) hand was exposed to ambient room temperature [mean (standard deviation; SD) temperature: 26.4 (0.5)°C, and humidity: 29 (7)%]. All trials commenced with a 10-min baseline phase, followed by a 1-h pre-immersion (Pre-CWI) phase. During both phases of all three trials, the left hand (up to ∼15 cm above the wrist) was placed in a custom-made, water-perfused, tube-lined mitten, and warm water was circulated maintaining the skin temperature of the fingers at ∼35.5°C (40). The baseline phase was void of any external (i.e., video and audio) stimulation. During the Pre-CWI phase of the cognitive+cold and control trials, subjects were asked to watch the documentary film “Our Planet” (Netflix, 2019). During the Pre-CWI phase of the cognitive→cold trial, subjects were requested to repeatedly perform a cognitive task battery via a computer-based platform [the Spaceflight Cognitive Assessment Tool for Windows (WinSCAT), NASA, USA; *see below for details*]. At the end of the Pre-CWI phase of all trials, the left hand was removed passively from the mitten, was covered with a thin plastic bag, and was immersed up to the ulnar and radial styloid for 30 min in a tank filled with 8°C stirring water (H-CWI phase). After the completion of the H-CWI phase, the hand was removed passively from the water, and a 15-min spontaneous hand rewarming (H-RW) phase ensued, during which the hand remained idle on the hand-support. During the H-CWI and H-RW phases of the cognitive→cold and control trials, subjects watched the documentary movie, whereas in these two respective phases of the cognitive+cold trial, they repetitively performed the cognitive task battery; the task was initiated 1 min before the start of the H-CWI phase.

**FIGURE 1.**
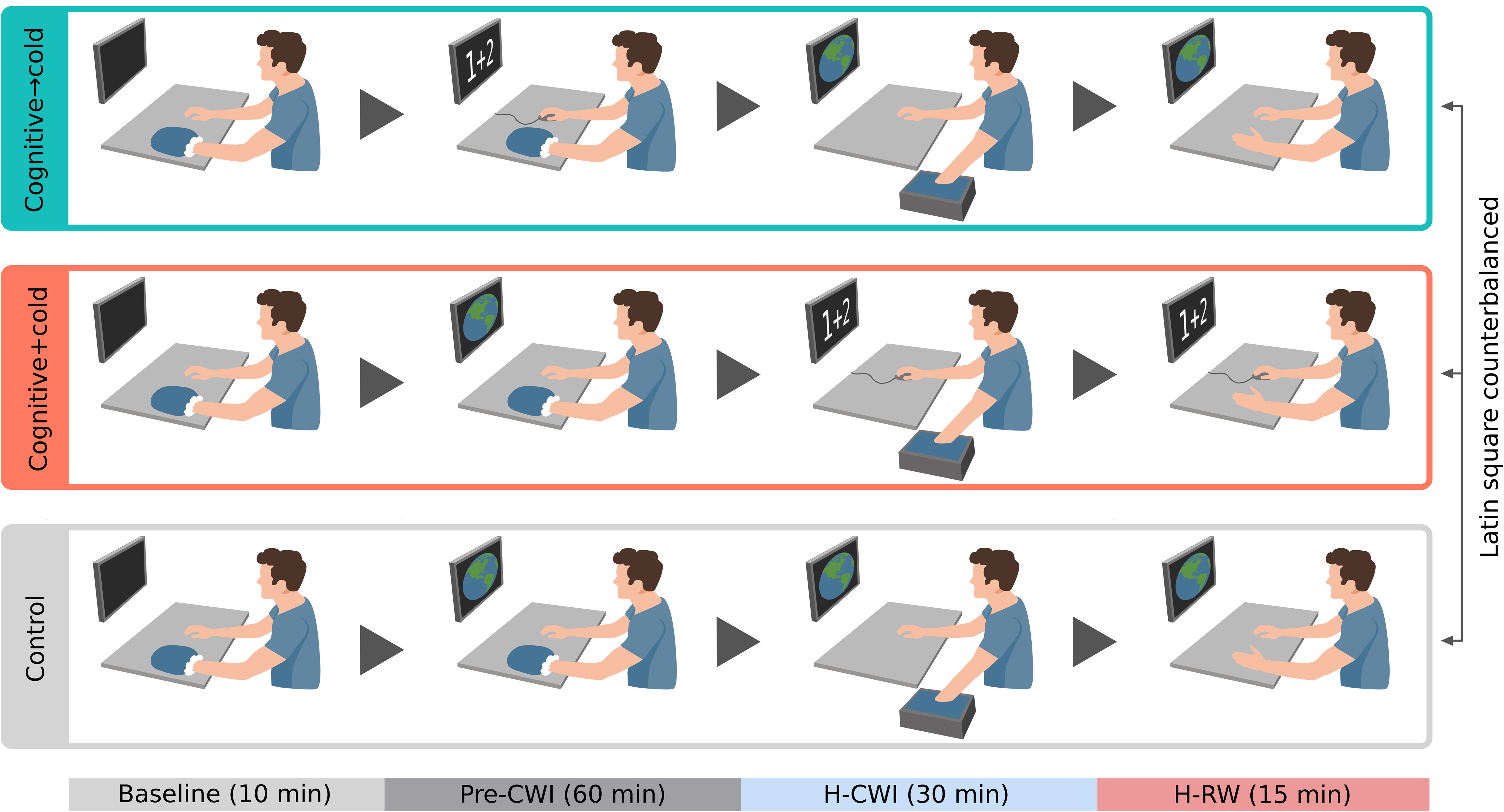
Schematic illustration of the testing protocol. Pre-CWI, a 60-min pre-cold water immersion phase; H-CWI: a 30-min hand cold water immersion phase; H-RW, a 15-min hand rewarming phase

Each cognitive task battery encompassed five executive-function subtests: code substitution, running memory, mathematical processing, match to sample, and delayed code substitution (41, 42). Subjects were requested to perform the cognitive task battery as many times as possible within the allocated phase of each trial. The interval between each consecutive task battery was ≤60-sec. No verbal feedback, or encouragement/harassment was provided during the task. Pilot trials in our laboratory indicated that repeated execution of the task required moderate levels of mental effort, leading to some degree of perceived fatigue. By contrast, the selection of the documentary “Our planet” was based on its content, which appears to be engaging, but emotionally neutral; it has been used to create an emotionally neutral condition in other mental-stress/fatigue studies (for instance, see Refs. 43-45).

The study was conducted between March and May. For the individual subject, the three experimental sessions were performed within a 2-week period, and always at the same time of the day, i.e., either in the morning (∼08:00; n = 4), or in the early (∼13:00; n = 6) or late (∼16:00; n = 2) afternoon. Subjects were always clad in T-shirt, shorts, and socks. During all trials, subjects were euglycemic [mean (SD) blood glucose: 5.3 (0.4) mmol·L^-1^; Accu-Check, Aviva, Roche, Manheim, Germany] and euhydrated (urine specific gravity <1.025; PAL-105 refractometer, ATAGO, Tokyo, Japan); both measures were conducted upon subjects’ arrival to the laboratory.

### Measurements and Instrumentation

#### Thermometry

All temperatures were sampled at 1 Hz with a NI USB-6215 data acquisition system, and processed with LabVIEW software (v. 2019, National Instruments, Austin, TX). The temperature of each finger of the left hand was continuously monitored with copper-constantan (T-type) thermocouple probes (Physitemp Instruments Inc, USA), attached to the middle of the palmar side of the distal phalanx. The average (*T*_F-avg_), minimum and maximum finger temperatures were then calculated. The total area under the curve (AUC) for *T*_F-avg_ was calculated for the H-CWI and H-RW phases, using the trapezoidal rule (GraphPad Software Inc., San Diego, CA). Furthermore, a finger skin temperature elevation of ≥1°C lasting for ≥3 min during the H-CWI phase was defined as a CIVD event. The following CIVD-related parameters were then assessed, using a custom-made program based on TestPoint (v7, Norton, MA): (i) the total number of CIVD events, (ii) the onset time of the first CIVD event, (iii) the temperature amplitude, i.e., the difference between the lowest temperature recorded just before the CIVD and the highest temperature reached during the CIVD, (iv) the duration of each CIVD event, and (v) the time interval between the lowest and the highest temperature of the CIVD.

Four additional thermocouples were placed on the upper arm, chest, thigh and calf, and the mean skin temperature was calculated. Rectal temperature was monitored with a thermistor probe (Yellow Springs Instruments, USA), inserted ∼10 cm beyond the anal sphincter.

#### Skin blood flux

Local skin blood flux was measured continuously, at a rate of 1 Hz, by laser-Doppler flowmetry (VMS-LDF2; Moor Instruments, Axminster, UK), using an optic probe (VP1/7; Moor Instruments, UK) that was placed on the palmar side of the distal phalanx of the left index finger, and the dorsal side of the right (i.e., non-immersed) forearm. The probes were positioned always by the same investigator, who employed anatomical landmarks to ensure that they were approximately at the same spots in all sessions. Before each session, the probes were calibrated against Brownian motion with a standardized colloidal suspension of polystyrene microspheres. Data were reported as cutaneous vascular conductance (CVC), which was calculated by dividing blood flux by mean arterial pressure (MAP). The AUC for finger CVC for the entire H-CWI and the last 10 min of the H-RW phases were also determined (GraphPad Software Inc., CA).

#### Arterial pressures and heart rate

Beat-to-beat systolic (SAP) and diastolic (DAP) arterial pressures, and MAP were measured continuously by finger photoplethysmography (Finometer, Finapress Medical Systems BV, the Netherlands), with the pressure cuff placed on the ring finger of the right hand, and with the reference pressure transducer positioned at the level of the heart. Heart rate was derived from the arterial pressure curve as the inverse of the interbeat interval. Cardiac stroke volume was estimated by a three-element model of arterial input impedance from the arterial-pressure waveform (Modelflow, Finometer; 46), and cardiac output was calculated by multiplying heart rate by stroke volume.

#### Cerebral oxygenation

Cerebral oxygenation was monitored continuously, at a rate of 1 Hz, by near-infrared spectroscopy (NIRS; NIRO-200NX, Hamamatsu Photonics, Japan). The probe was placed, at a fixed 4-cm interoptode distance, over the left prefrontal cortex at the midpoint between the Fp1 and F3 landmarks of the international electroencephalogram 10-20 system. Changes in oxygenated (Δ[O_2_Hb]) and deoxygenated (Δ[HHb]) hemoglobin concentration were determined; and total hemoglobin (Δ[tHb]) was derived from the sum of Δ[O2Hb] and Δ[HHb]. Data were expressed relative to the 10-min baseline phase.

#### Respiratory measurements

In all trials, subjects breathed through an oronasal mask, and expired ventilation (V̇e) and partial pressure of end-tidal carbon dioxide (PET_CO2_) were measured continuously with a metabolic unit (Metamax, 3B, Cortex Biophysik GmbH, Germany).

#### Perceptual measurements

During the baseline, Pre-CWI (at 15-min intervals), H-CWI (at minutes 1, 5, 10, 20, and 30), and H-RW (at minutes 5 and 15) phases, subjects were asked to provide ratings of their left-hand thermal sensation (from 1-cold to 7-hot), comfort (from 1-comfortable to 4-very uncomfortable), and pain (from 0-no pain to 10-unbearable pain). At the same time points, the perceived levels of mental effort (from 0-no mental effort to 10-very very high mental effort) and fatigue (from 1-not mentally fatigued to 5-mentally exhausted) were also assessed.

### Statistical analyses

All physiological data were reduced to 60-s averages. Normality of distribution was assessed using the Shapiro–Wilk test. Considering the study’s hypotheses (i.e., to evaluate differences from the control trial) and that the total amount of cognitive load between the cognition→cold and cognition+cold trials was unequal, the analysis was focused on the outcome of pairwise comparisons, wherein the responses obtained in the two intervention (i.e., cognition→cold and cognition+cold) trials were evaluated separately against those obtained in the control trial. Accordingly, temperature, cardiorespiratory and whole-body perceptual (i.e., mental effort and fatigue) responses were analyzed with two-way (trial × time) repeated measures analysis of variance (ANOVA). Sphericity was assessed using Mauchly’s test, and the Greenhouse–Geiser ε correction was applied when necessary. When ANOVA revealed significant effects, multiple pairwise comparisons were performed with Tukey’s honestly significant difference post hoc test. To account for unequal sample sizes comparisons (47), the NIRS data (n =11 for the cognition+cold trial, due to technical failure) and the cognitive performance output (for the exact n of the completed task-attempts, see the *results* section) were evaluated with linear mixed-effect models, with trial and time as fixed effects, and subjects as a random effect. The best fitting covariance structure was assessed with the corrected Akaike Information Criterion. Post hoc comparisons were made with Tukey’s honestly significant difference test for the NIRS data, and with the Dunnett’s test (i.e., differences from the 1^st^ attempt) for the cognitive performance data. Differences in AUCs, the regional perceptual (i.e., thermal sensation, discomfort and pain) responses and the CIVD parameters were assessed with Students’ paired two-tailed *t* test. Differences in the CIVD events were determined with the chi-square test. The data visualization and statistical analyses were performed using SPSS (IBM Corp. Released 2020. IBM SPSS Statistics for Windows, version 27.0. Armonk, NY: IBM Corp.) and Prism 9.0 (GraphPad Software Inc., San Diego, CA). Unless otherwise stated, data are presented as mean values with (SD). The *α* level of significance was set a priori at 0.05. Exact *P* values for single comparisons are reported down to *P* = 0.001; smaller values are reported as *P* < 0.001. The signs ≥ and ≤ are used to denote the smallest/biggest *P* value of several *P* values.

## RESULTS

The mean duration of each cognitive task battery was 14 (1) min. The median number of the completed cognitive-task attempts was 4 in the cognitive→cold trial (4 attempts: n = 9, and 3 attempts: n = 3), and 3 in the cognitive+cold trial (3 attempts: n = 10, and 2 attempts: n = 2). The overall cognitive performance was reduced gradually in both trials [by ∼16% in the cognitive→cold trial, and by ∼10% in the cognitive+cold trial; *P* ≤ 0.01; **FIGURE 2**). The cognitive task transiently enhanced the perceived levels of mental effort in both trials (*P* ≤ 0.03), as well as of mental fatigue, especially in the cognitive→cold trial (*P* < 0.01) (**FIGURE 3**).

**FIGURE 2.**
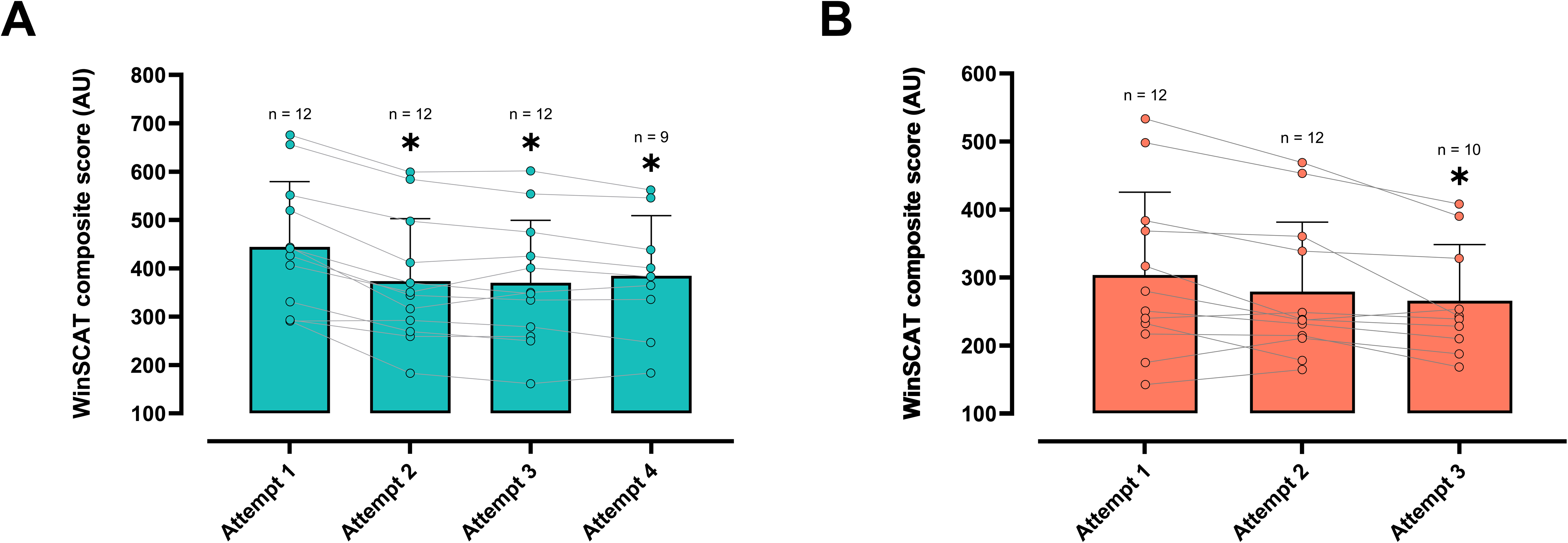
Mean (SD) and individual values of the composite score derived from the cognitive task battery [Spaceflight Cognitive Assessment Tool for Windows (WinSCAT)], performed repeatedly before (***A***; cognition→cold trial) and during (***B***; cognition+cold trial) the hand cold provocation (i.e., a 30-min immersion in 8°C water, followed by a 15-min rewarming). Data were analyzed with linear mixed-effects models with best-fit covariance structure, followed by Dunnett’s *post hoc* testing (*P* < 0.05). * Significantly different from the first attempt.

**FIGURE 3.**
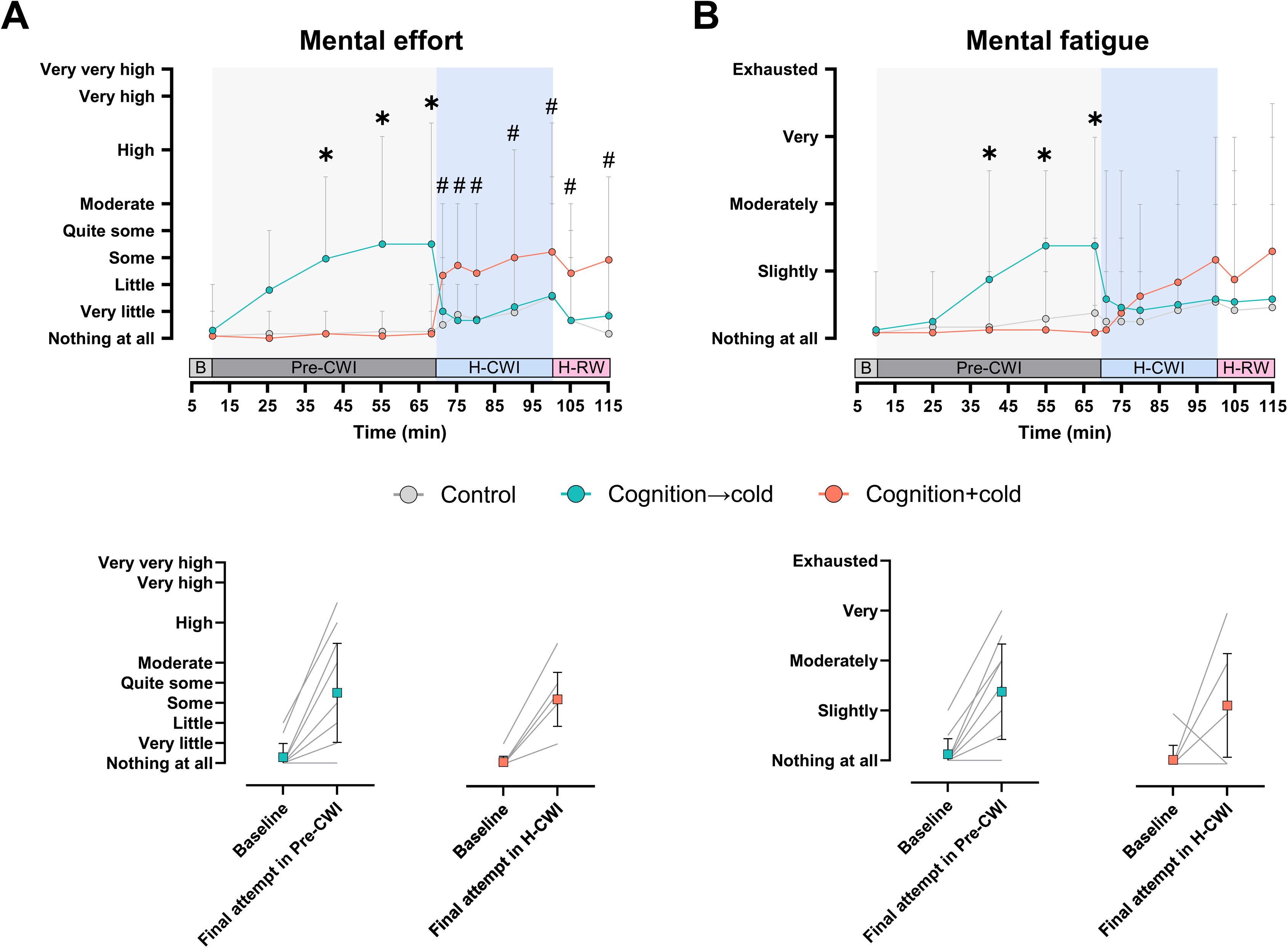
Mean (SD) values of the perceived levels of mental effort (***A***) and fatigue (***B***) obtained during the local cold provocation trials. Control trial, the hand cold provocation was performed without any cognitive task; Cognition→cold trial, the hand cold provocation was preceded by a 1-h cognitive task; Cognition+cold trial, the hand cold provocation was performed concomitantly with the cognitive task; B, a 10-min baseline phase; Pre-CWI, a 60-min pre-cold water immersion phase; H-CWI, a 30-min hand cold water immersion phase; H-RW, a 15-min hand rewarming phase. The *bottom* graphs depict the individual values during baseline and after the 4^th^ (in cognition→cold trial), and the 2^nd^ (in cognition+cold trial) attempt of performing the cognitive task battery. n = 12 healthy men. Data were analyzed with a 2-way repeated measured ANOVA, followed by Tukey’s honestly significant difference *post hoc* testing (*P* < 0.05). Significant difference * between the control and cognition→cold trials, and # between the control and cognition+cold trials.

### Whole-body responses

The whole-body physiological responses obtained during the three trials are summarized in **TABLE 2**. In all trials, subjects were euthermic, and their rectal and skin temperatures remained relatively constant (*P* ≥ 0.06). Stroke volume, cardiac output, V̇e, PET_CO2_ and forearm CVC did not vary across the trials (*P* ≥ 0.13). Furthermore, the absolute values of heart rate and arterial pressures were similar across all trials (*P* ≥ 0.19). Yet, the cognitive task enhanced the relative values (as calculated from the respective baseline phase) of heart rate and mean arterial pressure in the cognitive→cold trial (*P* < 0.03), and of heart rate in the cognitive+cold trial (*P* = 0.03). Moreover, compared to in the control trial, the magnitude of the cold-evoked MAP elevation (as calculated from the respective Pre-CWI phase), was attenuated in the cognitive→cold trial (*P* < 0.01), whereas, conversely, it was aggravated in the cognitive+cold trial (*P* = 0.03) (**FIGURE 4**). The cognitive task enhanced the cerebral Δ[tHb]: it increased, on average, by ∼405% in the cognitive→cold Pre-CWI phase (*P* = 0.03), and by ∼282% in the cognitive+cold H-CWI and H-RW phases (*P* ≤ 0.04). Δ[O_2_Hb] and Δ[HHb] did not vary significantly during the course of any of the three trials (*P* ≥ 0.45).

**FIGURE 4.**
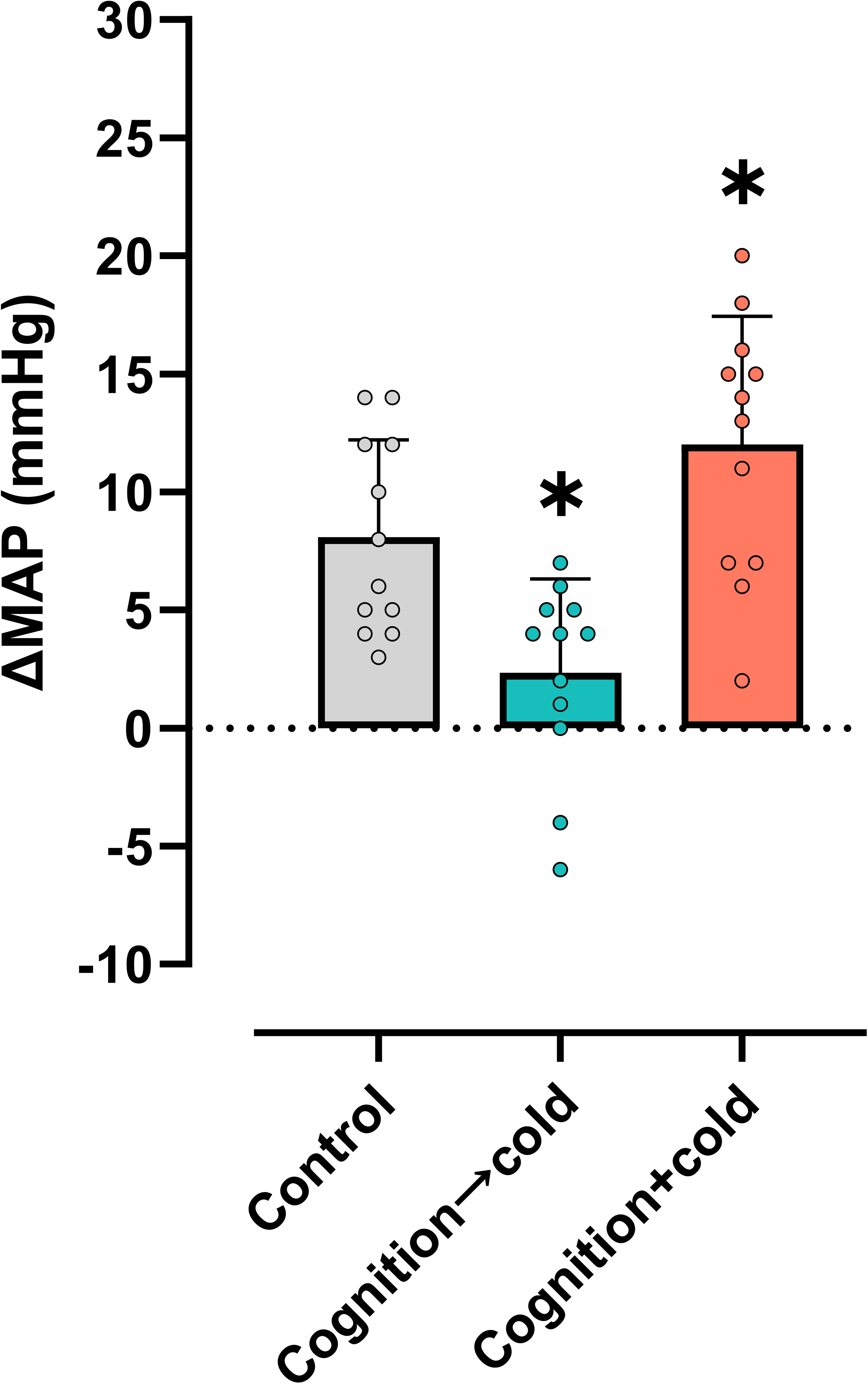
Mean (SD) and individual changes in mean arterial pressure (ΔMAP) from the pre-cold water immersion phase obtained during the 30-min hand cold-water immersion phase of the local provocation trials. Control trial, the hand cold provocation was performed without any cognitive task; Cognition→cold trial, the hand cold provocation was preceded by a 1-h cognitive task; Cognition+cold trial, the hand cold provocation was performed concomitantly with the cognitive task. n = 12 healthy men. Differences from the control trial were assessed with Student’s paired two-tailed *t*-tests (*P* < 0.05). * Significantly different from the control trial.

**TABLE 2.** Whole-body physiological responses obtained during the three hand cold provocation trials.

|  | Control trial |  |  |  | Cognition→cold trial |  |  |  | Cognition+cold trial |  |  |  |
| --- | --- | --- | --- | --- | --- | --- | --- | --- | --- | --- | --- | --- |
|  | Baseline | Pre-CWI | H-CWI | H-RW | Baseline | Pre-CWI | H-CWI | H-RW | Baseline | Pre-CWI | H-CWI | H-RW |
| Rectal temperature, °C | 37.1 (0.3) | 37.0 (0.2) | 37.0 (0.1) | 37.0 (0.1) | 37.1 (0.3) | 37.0 (0.2) | 37.0 (0.3) | 37.0 (0.3) | 37.2 (0.3) | 37.1 (0.2) | 37.0 (0.2) | 37.0 (0.2) |
| Skin temperature, °C | 32.1 (0.4) | 32.1 (0.4) | 32.1 (0.5) | 32.1 (0.6) | 32.3 (0.5) | 32.4 (0.6) | 32.3 (0.5) | 32.3 (0.5) | 32.2 (0.4) | 32.5 (0.4) | 32.7 (0.5) | 32.7 (0.5) |
| Heart rate, beats·min<sup>-1</sup> | 70 (5) | 70 (4) | 69 (9) | 70 (7) | 71 (8) | 76 (8) | 71 (9) | 71 (9) | 70 (5) | 70 (4) | 73 (7) | 73 (9) |
| ΔHeart rate, beats·min<sup>-1</sup> | 0 | 0 (3) | -1 (3) | 0 (4) | 0 | 3 (5)* | -2 (5) | -2 (5) | 0 | -1 (3) | 3 (5)* | 2 (5) |
| SAP, mmHg | 121 (8) | 121 (6) | 136 (7) | 134 (6) | 121 (11) | 126 (13) | 132 (10) | 131 (10) | 116 (10) | 115 (8) | 135 (11) | 135 (6) |
| ΔSAP, mmHg | 0 | 0 (8) | 15 (11) | 14 (11) | 0 | 5 (8) | 10 (9) | 9 (13) | 0 | -1 (6) | 18 (8) | 18 (7) |
| DAP, mmHg | 75 (7) | 76 (5) | 84 (7) | 83 (6) | 77 (6) | 80 (6) | 82 (6) | 84 (6) | 73 (6) | 73 (7) | 84 (6) | 83 (6) |
| ΔDAP, mmHg | 0 | 0 (4) | 9 (5) | 8 (6) | 0 | 3 (4) | 5 (4) | 5 (6) | 0 | 0 (3) | 11 (3) | 11 (4) |
| MAP, mmHg | 92 (7) | 92 (6) | 103 (7) | 101 (7) | 93 (8) | 97 (8) | 100 (8) | 103 (7) | 89 (7) | 88 (7) | 103 (7) | 101 (7) |
| ΔMAP, mmHg | 0 | 0 (5) | 11 (7) | 9 (8) | 0 | 4 (5)* | 7 (6) | 6 (9) | 0 | -1 (4) | 14 (5) | 13 (5) |
| Stroke volume, ml | 84 (11) | 83 (10) | 83 (15) | 84 (17) | 86 (17) | 87 (18) | 89 (17) | 89 (12) | 84 (19) | 82 (16) | 86 (19) | 85 (19) |
| Cardiac output, L·min<sup>-1</sup> | 5.8 (0.9) | 5.7 (0.9) | 5.6 (1.1) | 5.8 (1.3) | 6.5 (1.3) | 6.4 (1.5) | 6.4 (1.4) | 6.2 (1.0) | 5.8 (1.1) | 5.7 (1.1) | 6.3 (1.4) | 6.1 (1.5) |
| $\dot{V}_E$, L·min<sup>-1</sup> | 10.3 (1.4) | 10.5 (1.5) | 11.4 (1.7) | 11.9 (2.0) | 11.2 (1.5) | 11.2 (1.3) | 11.3 (1.5) | 11.2 (1.3) | 11.0 (2.0) | 11.2 (1.7) | 11.8 (1.7) | 11.9 (2.0) |
| PETco<sub>2</sub>, mmHg | 36.1 (3.0) | 35.7 (2.2) | 35.1 (2.3) | 34.6 (3.5) | 35.5 (2.4) | 35.5 (2.1) | 35.6 (2.3) | 35.1 (2.3) | 35.7 (3.5) | 35.6 (3.1) | 35.2 (1.8) | 34.9 (2.1) |
| Forearm CVC, PU·mmHg<sup>-1</sup> | 0.6 (0.3) | 0.6 (0.3) | 0.4 (0.1) | 0.4 (0.1) | 0.5 (0.3) | 0.5 (0.1) | 0.4 (0.2) | 0.4 (0.2) | 0.6 (0.3) | 0.6 (0.3) | 0.5 (0.2) | 0.5 (0.2) |
| Δ[O<sub>2</sub>Hb], μM | 0 | 0.1 (2.7) | 0.5 (3.9) | -1.0 (3.8) | 0 | 0.9 (2.3) | 0.3 (3.8) | -0.3 (3.8) | 0 | 0.5 (1.9) | 1.8 (1.6) | -1.0 (3.8) |
| Δ[HHb], μM | 0 | 0.0 (1.5) | -0.4 (2.4) | -0.3 (2.6) | 0 | -0.7 (1.4) | -0.5 (2.3) | -0.4 (1.8) | 0 | -0.1 (0.7) | -0.8 (1.2) | -0.3 (2.6) |
| Δ[tHb], μM | 0 | -0.4 (1.5) | 0.1 (1.9) | -1.0 (1.8) | 0 | 1.3 (1.6)* | 0.8 (1.8) | 0.4 (1.9) | 0 | -0.1 (1.1) | 1.5 (1.6)* | -1.0 (1.8)* |

Values are mean (SD). Systolic (SAP), diastolic (DAP) and mean (MAP) arterial pressures, expired ventilation ( $\dot{V}_E$ ), partial pressure of end-tidal carbon dioxide (PETco<sub>2</sub>), cutaneous vascular conductance (CVC), changes in oxygenated (Δ[O<sub>2</sub>Hb]), deoxygenated (Δ[HHb]), and total (Δ[tHb]) hemoglobin. n = 12 healthy men, aside from Δ[O<sub>2</sub>Hb], Δ[HHb] Δ[tHb] where n = 11 in the Cognition+cold trial. Control trial, the hand cold provocation was performed without any cognitive task; Cognition→cold trial, the hand cold provocation was preceded by a 1-h cognitive task; Cognition+cold trial, the hand cold provocation was performed concomitantly with the cognitive task; Baseline, a 10-
1 min baseline phase; Pre-CWI, a 60-min pre-cold water immersion phase; H-CWI: a 30-min hand cold water immersion phase; H-RW, a 15-min hand rewarming phase. Data 2 were analyzed with 2-way repeated measured ANOVA, except for $\Delta[\text{O}_2\text{Hb}]$ , $\Delta[\text{HHb}]$ $\Delta[\text{tHb}]$ for which data were analyzed with linear mixed-effects models with best-fit 3 covariance structure. Both models were followed by Tukey's honestly significant difference *post hoc* testing. ( $P < 0.05$ ). \* Significantly different from the control trial.

### Finger responses

#### Cognitive→cold vs. control trials

During the baseline and Pre-CWI phases, neither *T*_F-avg_ nor finger CVC differed between trials (*P ≥* 0.20; **FIGURE 5A** and **D**). During the H-CWI and H-RW phases, *T*_F-avg_ was similar in the two trials (*P* ≥ 0.62; **FIGURE 5A, B** and **C**). No intertrial differences were noted in the minimum and maximum temperatures, nor the incidence and the temperature and temporal characteristics of the CIVD events (*P* ≥ 0.07; **TABLE 3**). The cold-induced drop in finger CVC, however, was blunted during the first 9 min of the cognitive→cold H-CWI phase (*P* < 0.001; **FIGURE 5D**). AUC for finger CVC was also higher in the cognitive→cold than the control H-CWI phase (*P* = 0.02; **FIGURE 5E**), whereas it did not differ in the H-RW phase (*P* = 0.20; **FIGURE 5F**). During the cognitive→cold H-CWI phase, the thermal discomfort was attenuated (*P* = 0.05), but not the sensations of coldness (*P* = 0.70) and pain (*P* = 0.07) (**FIGURE 6**).

**FIGURE 5.**
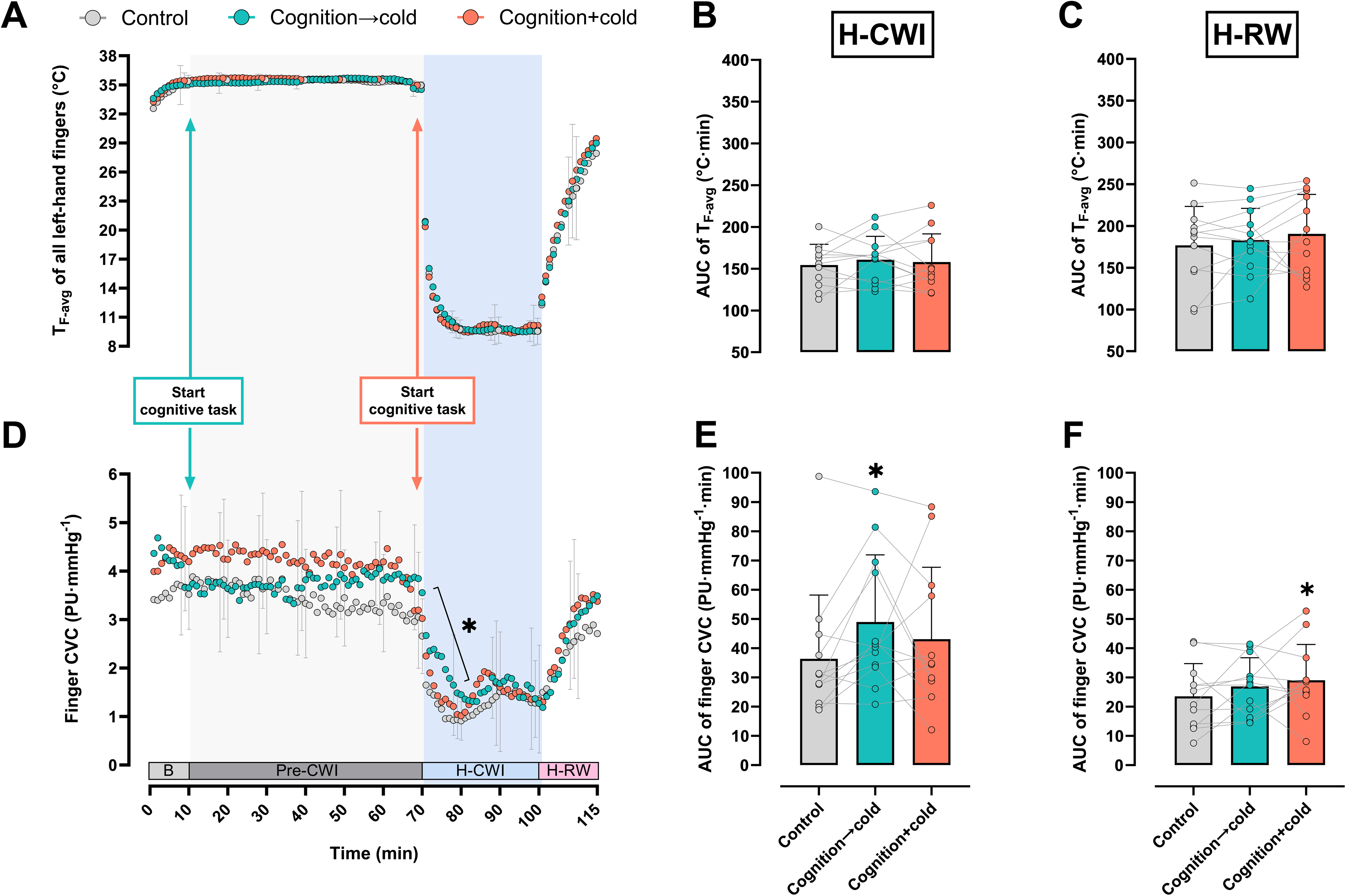
Time course [mean (SD)] of skin temperature (*T*_F-avg_) of all five fingers (***A***), and cutaneous vascular conductance (CVC) of the index finger (***B***) of the left hand during the local cold provocation trials. Mean (SD) values of the area under the curve (AUC) for *T*_F-avg_ (***C***) and finger CVC (***D***) determined during the hand cold-water immersion and rewarming phases. Control trial, the hand cold provocation was performed without any cognitive task; Cognition→cold trial, the hand cold provocation was preceded by a 1-h cognitive task; Cognition+cold trial, the hand cold provocation was performed concomitantly with the cognitive task; B, a 10-min baseline phase; Pre-CWI, a 60-min pre-cold water immersion phase; H-CWI: a 30-min hand cold water immersion phase; H-RW, a 15-min hand rewarming phase. n = 12 healthy men. The absolute *T*F-avg and finger CVC values were analyzed with 2-way repeated measured ANOVA, followed by Tukey’s honestly significant difference *post hoc* testing. The AUC values for *T*_F-avg_ and finger CVC were analyzed with Student’s paired two-tailed *t*-tests. (*P* < 0.05). * Significantly different from the control trial.

**FIGURE 6.**
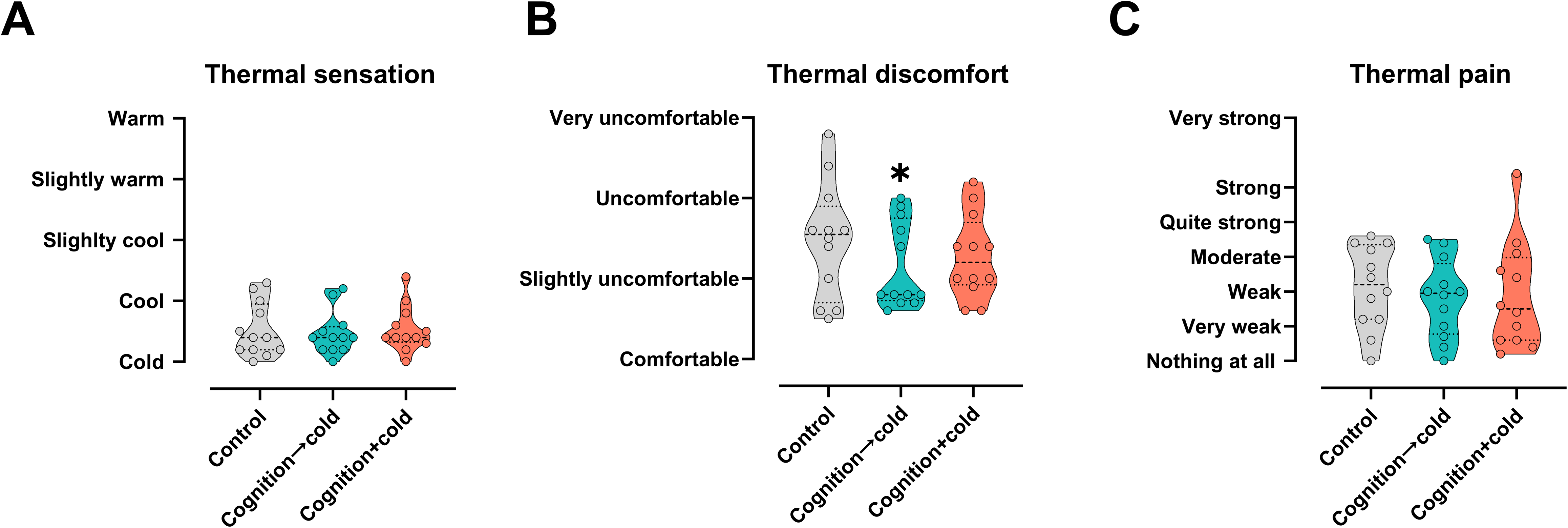
Violin plots of thermal sensation (***A***), discomfort (***B***), and pain (***C***) obtained during the 30-min hand cold-water immersion phase of the local cold-provocation trials. Control trial, the hand cold provocation was performed without any cognitive task; Cognition→cold trial, the hand cold provocation was preceded by a 1-h cognitive task; Cognition+cold trial, the hand cold provocation was performed concomitantly with the cognitive task. n = 12 healthy men. Differences from the control trial were assessed with Student’s paired two-tailed *t*-tests (*P* < 0.05). * Significantly different from the control trial.

**TABLE 3.** Minimum and maximum temperature, number of cold-induced vasodilation events, and temperature amplitude, duration, time onset and time interval of cold-induced vasodilation on the palmar side of the distal phalanx of the left-hand fingers during the 30-min hand cold water immersion phase of the three provocation trials.

|  | Control trial | Cognition→cold trial | Cognition+cold trial |
| --- | --- | --- | --- |
| Minimum temperature, °C | 8.7 (0.5) | 8.8 (0.5) | 8.7 (0.6) |
| Maximum temperature, °C | 10.8 (2.0) | 10.9 (1.4) | 11.1 (2.3) |
| CIVD events, no. | 4 (4) | 4 (3) | 5 (5) |
| CIVD amplitude, °C | 2.7 (1.8) | 2.4 (0.7) | 2.8 (1.0) |
| CIVD duration, min | 7.9 (2.0) | 6.3 (1.7) | 6.9 (2.1) |
| CIVD onset time, sec | 656 (195) | 754 (313) | 819 (362) |
| CIVD time-interval, sec | 283 (81) | 216 (63) | 274 (87) |

Values are mean (SD). CIVD, cold induced vasodilation. Control trial, the hand cold provocation was performed without any cognitive task; Cognition→cold trial, the hand cold provocation was preceded by a 1-h cognitive task; Cognition+cold trial, the hand cold provocation was performed concomitantly with the cognitive task. n = 12 healthy men. Differences from the control trial were assessed with Student's paired t-tests, except for the number of CIVD events that were assessed with the chi-square test ( $P > 0.05$ ).

#### Cognitive+cold vs. control trials

Regardless of the phase, no intertrial differences were observed in *T*_F-avg_ and finger CVC (*P* ≥ 0.35; **FIGURE 5**). Yet AUC for finger CVC was higher in the cognitive+cold than the control H-RW phase (*P* = 0.05; **FIGURE 5F**). The minimum and maximum temperatures, as well as the incidence and the characteristics of the CIVD events were similar in the two trials (*P* ≥ 0.28; **TABLE 3**). The sensation of coldness, thermal discomfort, and pain did not vary between the two trials (*P* ≥ 0.28; **FIGURE 6**).

## DISCUSSION

Cutaneous vasomotor tone is governed primarily by changes in body core and/or surface temperature, but is also modulated by non-thermal factors (1, 2). The present study hence aimed to examine acral-skin vasomotor and thermoperceptual responsiveness to localized cooling during two discrete experimental paradigms of enhanced cognitive strain. Finger temperature, circulatory and perceptual responses were thus monitored during and after a 30-min hand immersion in 8°C stirring water, performed either immediately after a 60-min continual execution of a cognitive task battery, or during the simultaneous performance of the cognitive task. Subjects’ autonomic and perceptual responses were compared with those obtained in a control cold-provocation trial, wherein they watched an emotionally-neutral documentary. All experimental sessions were conducted in identical laboratory settings, which were carefully controlled with regards to external arousal stimuli (e.g., noise, feedback), subjects’ whole-body temperature and posture, as well as the pre-immersion temperature of the tested hand. The main findings of the study were that: (i) the preceding execution of a prolonged cognitive stress task leading to moderate levels of mental fatigue, temporary attenuated finger vasoconstriction and alleviated thermal discomfort during local cooling, and (ii) the concurent administration of cognitive and cold stress did not modify finger vasoreactivity or thermosensitivity during cooling, but accelerated finger reperfusion following cooling.

### Independent effects of sustained cognitive loading

As anticipated, the employed cognitive task battery, which comprised a variety of executive functioning components (42), required, on average, moderate levels of sustained cognitive effort. The subjective assessment of mental exertion was also accompanied by a consistent elevation of prefrontal cortex oxygenation, presumably reflecting a regional increase in neural activity and/or in blood flow (48, 49). Although, to avoid provoking profound emotional distress, we did not impose any specific pressure on the duration and the number of task attempts, the extended period of cognitive effort invariably resulted in progressive mental fatigue and gradually impaired task performance (i.e., a “time-on-task” effect; 50, 51). Moreover, as commonly during cognitive-stress interventions (52–55), a state of mild sympathetic hyperactivity transpired during the cognitive-loading period, as indicated by the small but persistent elevations in heart rate and mean arterial pressure.

In the cognition→cold trial, the initiation of the cognitive task caused an instant drop in finger CVC (by ∼7%; *P* = 0.07), confirming the high sensitivity of the acral-skin vasculature to arousal stimuli (9, 34). Forearm CVC, by contrast, did not change in response to the cognitive task; a result that is somewhat at variance with the common finding of slightly increased blood flow in non-acral skin during acute mental stress (16–19, 21). Given that the right hand was used for clicking the computer-mouse during the cognitive task, and that the forearm CVC measurements were conducted on the same arm, we cannot exclude that the forearm laser-Doppler recordings were confounded by small arm-muscle activity – although no excessive noise was noted in the raw CVC data. Notwithstanding, assuming similar responsiveness of the precapillary resistance vessels in acral and non-acral cutaneous beds, the present discrepancy in CVC response between fingertip and forearm probably reflected closing of finger AVAs in response to the cognitive task, since AVAs are abundant in the fingertip (7). Still, the finger constrictor response did not prevail during the entire cognitive task period, but abated at approximately minute 30 of the Pre-CWI phase. Notably, the restoration of finger CVC to its baseline values appeared to coincide with the augmented sensations of mental fatigue. Although speculative, the recovery of fingertip blood flow during the course of the cognitive task might be associated with accumulation of circulating adrenal-hormones (e.g., adrenaline; 53, 56, 57) and/or release of metabolites (e.g., nitric oxide; 17), typically induced by mental strain, and hence contributing to the vasodilatory response of the cutaneous arterioles (17, 19, 58).

### Combined effects of sustained cognitive loading & cold

A novel finding of this study was that, in the cognition→cold trial, the 60-min cognitive task blunted, at least initially, the magnitude of finger vasoconstriction instigated by the subsequent application of cold stress. The experimental design does obviously not allow us to determine the exact mechanism(s) underlying the response. Judging by the concurrent diminution of the systemic pressor reactivity to local cooling (**FIGURE 4**), we speculate that the transient attenuation of finger vasoreactivity to thermal stress probably reflected a peripheral adrenergic desenzitization, elicited by sustained elevation of circulating catecholamines secondary to the mental-fatigue state (cf. 59, 60). A similar autonomic hypoactivation to other sympathoexcitatory maneuvers (such as exercise, whole-body cooling) has been detected in mentally-fatigued individuals after a period of sleep deprivation (61, 62). That the acral-skin vasomotor behavior to cold was modified only in the cognition→cold trial, also suggests that the response was driven by the cumulative volume of cognitive loading and associated fatigue development, rather than by the acute bout of demanding cognitive activity. Indeed, contrary to in the cognition→cold trial, the constrictor response of finger arterioles was unaffected in the cognition+cold trial, wherein the cognitive task was initiated only 1 minute prior to the application of local cooling. The co-existence of the cold and cognitive stressors, in fact, produced an additive interaction (63), upregulating the arterial-pressure responsiveness to cold stimulation. Still, in the cognition+cold trial, the reperfusion of the fingertip was enhanced after the removal of the cold stressor and the continuation of the mental task, as indicated by the increased CVC values during the last 10 min of the H-RW phase. Again, the facilitation of the rewarming response might have been mediated by the vasodilatory effects of increased concentrations of hormones (e.g., adrenaline) and/or vasoactive substances (e.g., nitric oxide) resulting from mental fatigue (17, 19, 58). Of interest is that a somewhat similar rewarming pattern was displayed also in the cognitive→cold trial, albeit the difference was not statistically significant.

Neither the incidence nor the magnitude of CIVD response was affected by the intervention trials. This finding is in conflict with those in two previous investigations (29, 31). The discrepancy between the CIVD reponses in present and aforementioned studies can probably be attributed to the differences in the severity and application mode of the mental stressor. Thus, in the previous studies, the CIVD response was suppressed either in individuals suffering from long-term, intense anxiety (31), or when a strong emotional (unpleasant) stimulus was introduced suddenly at the temperature-zenith phase of a CIVD wave (29). Our results clearly show that moderate levels of sustained cognitive strain, imposed either before or during local cooling, do not exert any prominent influence on the finger CIVD response.

In the cognition→cold trial, the attenuation of finger cold-induced vasoconstriction was attended by a persistent alleviation of thermal discomfort; whereas subjects’ discriminative and noxious sensations to cold remained unchanged. It is plausible that the enhanced hedonic perception reflected a general improvement in subjects’ affective valence (64), engendered by shifting their focus from a repetitive and mentally demanding task to a physical task requiring less cognitive resources. Notably, the subjects’ perceptions of mental effort and fatigue dissipated immediately upon the cognitive-task termination and the subsequent hand immersion in cold water. Contrary to in the cognition→cold trial, superimposition of cognitive loading on the cold stimulus did not mofidy the perception of thermal discomfort. Apparently, in this condition, the 8°C cold water provoked strong unpleasant and noxius sensations, which overrode any potential distraction effects derived from the mental stress task (cf. 65, 66). Collectively, these findings suggest that, in conditions of moderately enhanced cognitive strain, the central processing of thermo-afferent information remains intact.

### Methodological considerations

During all Pre-CWI phases, the temperature of the fingers was clamped succesfully at ∼35.5°C. It might, however, be argued that the numerically higher values of finger CVC during the last part of the Pre-CWI phase in the cognitive→cold trial than that in the control trial, may have determined the temporary reduction of cold-induced vasoconstriction observed in the H-CWI phase (cf. Ref. 67). Such an explanation, however, seems less likely, because, in the cognitive+cold trial, in which the pre-immersion finger CVC was also somewhat enhanced, finger vasoreactivity to cold stimulus remained unaffected. We, nevertheless, acknowledge that the study would have benefited from a physiological calibration of the laser-Doppler recordings, e.g., the data normalization to a maximal CVC value produced by local heating application (68).

In line with previous works (69), the perceived magnitude of mental effort and fatigue during the cognitive task, described a large inter-individual variability, with some subjects being more affected than others. This response variation was independent of the subjects’ individual characteristics (e.g., age, chronotype). It has been suggested that the vascular reactions to a mental stress task might be associated with the subjective levels of perceived stress/fatigue (70). Yet, in the present study, we did not detect any correlation between the AUC for finger CVC and the perceived mental effort, in either of the intervention trials (cognitive→cold trial: *r* = 0.17, *P* = 0.60 and cognitive+cold trial: *r* = 0.34, *P* = 0.28).

Our observations are limited to the acral skin vasculature of the hand of young healthy men. Future work should evaluate whether the combined effects of cold and mental stress are limb and/or sex dependent; their separate examination suggests that between-limb and -sex variations to local cold (32, 71, 72) and mental stress (54, 73) may exist. Lastly, measurements of stress-hormones (e.g., epinephrine, norepinephrine) and vasoactive substances (e.g., nitric oxide) would have facilitated the interpetation of our results.

### Practical perspectives

Ostensibly, current findings may suggest that enhanced cognitive strain may reduce the risk of local cold injury. It should be noted, however, that the present investigation concerned autonomic and perceptual responses to slight-to-moderate levels of mental stress. Thus, although moderate levels of prolonged cognitive exertion do not seem to substantially affect peripheral hemodynamic and perceptual responses to local cold, it remains unknown if, or to what extent, severe mental stress/fatigue, likely to be encountered in real-life scenarios, compromises the capacity and efficiency of conscious thermobehavioral actions (i.e., our first line of cold defence). For instance, previous works have demonstrated that the ability to make favorable decisions in risky conditions may be impinged by cognitive overloading (74–76). Valenza et al. (77) have also found that the cold-induced impairment in complex hand-manipulation skills is exacerbated by mental fatigue provoked by a 35-min cognitive task.

### Conclusions

The present findings demonstrate that acral-skin vasomotor responsiveness to localized cooling may be modulated by sustained cognitive loading. Thus, when cold stimulation is applied in moderately mentally-fatigued individuals, finger cold-induced vasoconstriction is transiently attenuated, and thermal discomfort is mitigated. The concurrent application of enhanced loads of mental effort and local cooling does not potentiate finger vasoconstriction during cooling but appears to facilitate digit reperfusion following cooling. Further studies are required to investigate the mechanisms underlying the synergy of thermal and mental stressors on the function of acral-skin vasculature.

## DATA AVAILABILITY

Data will be made available upon reasonable request.

## ACKNOWLEDGEMENTS

We are grateful to all the subjects for participation. We acknowledge Ms. Dilja Sayfulaeva for valuable assistance.

## AUTHOR CONTRIBUTIONS

M.I.M., A.E., O.E. and M.E.K. conceived and designed research; M.I.M., A.E. and M.E.K. performed experiments; M.I.M. analyzed data; M.I.M. and M.E.K. interpreted results of experiments; M.I.M. prepared figures and drafted manuscript; M.I.M. and M.E.K. edited and revised manuscript; M.I.M., A.E., O.E. and M.E.K. approved final version of manuscript.

